# Nutrient limitation mimics artemisinin tolerance in malaria

**DOI:** 10.1101/2022.06.14.496121

**Authors:** Audrey Brown, Michelle D Warthan, Anush Aryal, Shiwei Liu, Jennifer L Guler

## Abstract

Mounting evidence demonstrates that nutritional environment can alter pathogen drug sensitivity. While the nutrient-rich media used for standard *in vitro* culture contains supra-physiological nutrient concentrations, pathogens encounter a relatively restrictive environment *in vivo*. We assessed the effect of nutrient limitation on the protozoan parasite that causes malaria and demonstrated that short-term growth under physiologically-relevant mild nutrient stress (or “metabolic priming”) triggers increased tolerance of the potent antimalarial drug dihydroartemisinin (DHA). We observed beneficial effects using both short-term survival assays and longer-term proliferation studies, where metabolic priming increases parasite survival to a level previously defined as DHA resistant (>1% survival). We performed these assessments by either decreasing single nutrients that have distinct roles in parasite metabolism or using a media formulation with reductions in many nutrients that simulates the human plasma environment. We determined that priming-induced DHA tolerance was restricted to parasites that had newly invaded the host red blood cell but the effect was not dependent on genetic background. The molecular mechanisms of this intrinsic effect mimic aspects of genetic artemisinin tolerance, including translational repression, autophagy, and protein export. This finding suggests regardless of the impact on survival rates, environmental stress could stimulate changes that ultimately directly contribute to drug tolerance. Because metabolic stress is likely to occur more frequently *in vivo* compared to the stable *in vitro* environment, priming-induced drug tolerance has ramifications for how *in vitro* results translate to *in vivo* studies. Improving our understanding of how pathogens adjust their metabolism to impact survival of current and future drugs is an important avenue of research to prevent and slow the spread of resistance.

## Introduction

*In vitro* culture provides microbes with a stable environment for growth with relatively little challenge to organismal homeostasis. For example, the supra-physiological concentrations of nutrients *in vitro* largely protect organisms from the stress of starvation. Yet, nutrient limitation occurs frequently *in vivo*^1-4^. Therefore, the consequences of dynamic adaptation to stressors, such as nutrient variation, are largely missed in standard *in vitro* experiments.

Across the tree of life, the relationship between stress and fitness consequences is not necessarily linear. The adaptive benefits of low doses of mild environmental stress overwhelm the adverse effects from the stressor in a biphasic dose response phenomenon termed hormesis^5,6^ (**Fig. 1A**). Given this effect, adjustment of metabolism in response to mild nutrient stress has the potential to increase pathogen fitness and survival of subsequent stressors, such as drug pressure.

**Figure 1.**
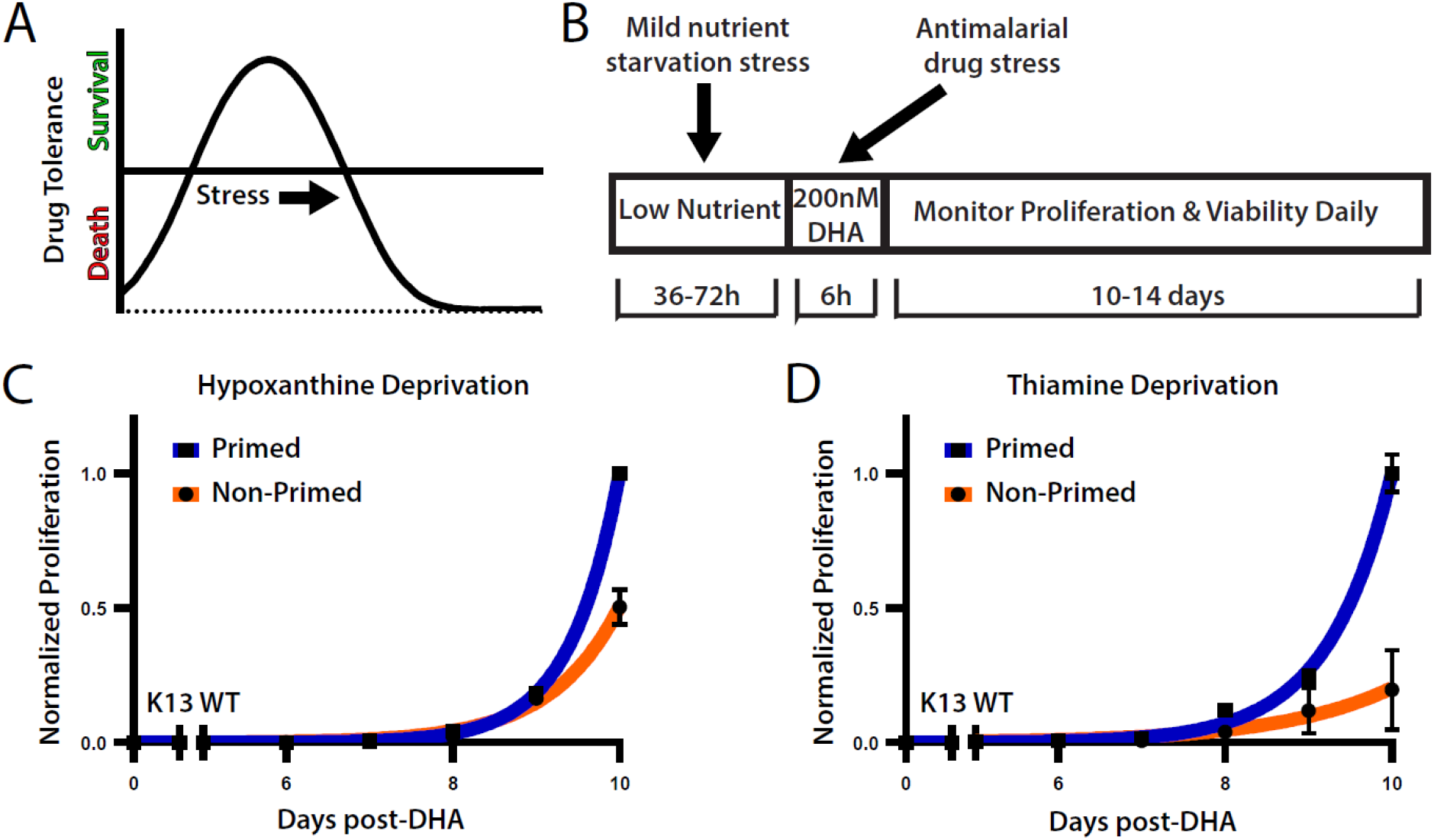
Mild nutrient stress prior to DHA treatment increases post-drug recovery. A) The biphasic dose response to stress characteristic of hormesis. B) Paradigm used for metabolic priming followed by DHA challenge and recovery. C-D) Post-DHA recovery in standard media of *P. falciparum* primed in either low hypoxanthine or thiamine-free media. Bars represent S.E.M. of technical replicates within one representative experiment. Results from all independent assays are detailed in Supplemental Table 1 (low hypoxanthine; *N=*5, thiamine-free; *N=*4).

Indeed, metabolic adjustments under varied nutritional environments alter drug sensitivity across diverse pathogens^7,8^. In *Trypanasoma cruzi*, reduction of exogenous glutamine decreases flux through the sterol synthesis pathway rendering parasites sensitive to azole drugs^9^. *Pseudomonas aeruginosa* supplementation with specific carbon sources increases susceptibility to tobramycin^10^. Increasing proline concentration confers halofuginone resistance in *Plasmodium falciparum*^11^. The nutritional environment *in vitro* is highly divergent from conditions that pathogens are exposed to in the context of human infection, with nutrient levels frequently found at considerably higher levels in culture^12-14^. Discrepancies between *in vitro* and *in vivo* environments can lead to reduced translatability of findings from non-physiological culture-based experiments when metabolic flux through pathways is altered^15,16^. For example, glycerol metabolism sensitizes *Mycobacterium tuberculosis* to pyrimidine-imidazoles through accumulation of toxic metabolites; however, these drugs are ineffective *in vivo* where changes in nutrient availability lead to differences in carbon metabolism^15^.

**Table 1.** Length and concentration of nutrient limitation for metabolic priming. *Priming was considered successful for this nutrient when an approximately 10-50% growth decrease occurred, there was little to no decrease in mitochondrial membrane potential (MMP) (approx. <10% drop), and ring stage percentages were within <10% ± non-primed. **Priming was considered successful for these nutrients when the latter two above conditions (MMP and staging) were met but no growth reduction criteria were required.

| Nutrient Restricted | Class of Restricted Nutrient | Conc. of Restricted Nutrient | Length of Restriction |
| --- | --- | --- | --- |
| Hypoxanthine* | Purine | 0.5uM | 48-72h |
| Thiamine** | Cofactor | 0.0uM | 36h |
| Multiple (HPLM)** | Multiple | Variable | ≥ 7d |

These previously characterized changes in pathogen drug sensitivity as a consequence of varied nutritional environment typically depend on either 1) changes in flux through a specific metabolic pathway relevant to a given drug target or 2) a general slowing of growth leading to decreased uptake and activation of drug. Here, we present a case where nutrient stress induces a change in drug sensitivity of the human malaria parasite, *P. falciparum*. We demonstrate that short-term growth under mild, physiologically relevant nutrient limitation leads to superior recovery from drug treatment. We observe this effect under distinct conditions that impact the parasites differently; therefore, it is not dependent on a change in flux within a specific metabolic pathway nor an overall decrease in growth rate (as in above examples). Importantly, the increased survival rates achieve a level that reaches the field’s definition of “resistance,” yet genetic mutations that confer resistance are not necessary. We determined that the induction of pro-survival pathways within the parasite occurs during nutrient limitation, which prime the parasite to better withstand drug treatment. This study emphasizes the link between nutrient limitation, pro-survival pathways, and sensitivity to drugs and may hold important implications for the evolution of drug resistant pathogens.

## Results

### Metabolic priming increases parasite recovery from DHA

We sought to determine the effect of mild nutrient stress on the survival of *P. falciparum*. Asynchronous parasite cultures were subjected to short-term (36-72h) incubation in media restricted of either hypoxanthine or thiamine (**Table 1**, termed “metabolic priming”). *Plasmodium* lack the ability to *de novo* synthesize purines, instead requiring scavenging from the environment. In serum-free malaria cultures, the purine derivative, hypoxanthine, is a standard additive to fulfill this requirement. In contrast, *Plasmodium* can both *de novo* synthesize or scavenge thiamine^17^. In addition to different modes of acquisition for these nutrients, hypoxanthine and thiamine represent distinct metabolite classes: a nucleoside important for nucleic acid synthesis and a cofactor for apicoplast and mitochondrial metabolism, respectively. The duration of incubation and level of nutrient limitation for these metabolites were optimized to induce mild stress while keeping the majority of the parasite population viable (**Supplemental Fig. 1**).

We treated parasites grown in either metabolic priming conditions or standard media (formulations detailed in *Methods*) with a rapid-acting, highly potent antimalarial drug (200nM dihydroartemisinin (DHA) for 6h, **Fig. 1B**). DHA is an artemisinin derivative that causes widespread damage within parasites by promiscuously alkylating different classes of biomolecules, including heme and numerous proteins, thereby disrupting multiple biological pathways^18-20^. Following removal of drug and continued culture for 10 days in standard media, primed parasites showed a significant increase in cumulative proliferation compared to non-primed controls (low hypoxanthine mean day 10 difference: 130%, *N=* 5, *p =* 0.0126; thiamine-free: 173%, *N=4, p =* 0.0074) (**Fig. 1C-D**). We detected this effect in both priming conditions despite the restricted nutrients in each condition being of unrelated metabolite classes, irrespective of the parasite line (**Supplemental Table 1**, *Dd2* originally from Southeast Asia and *NF54* from Africa). The increased cumulative proliferation is driven by altered response to DHA treatment; we only observed this effect in primed parasites that were subjected to DHA treatment but not in those treated with vehicle alone (**Supplemental Fig. 2**).

**Figure 2.**
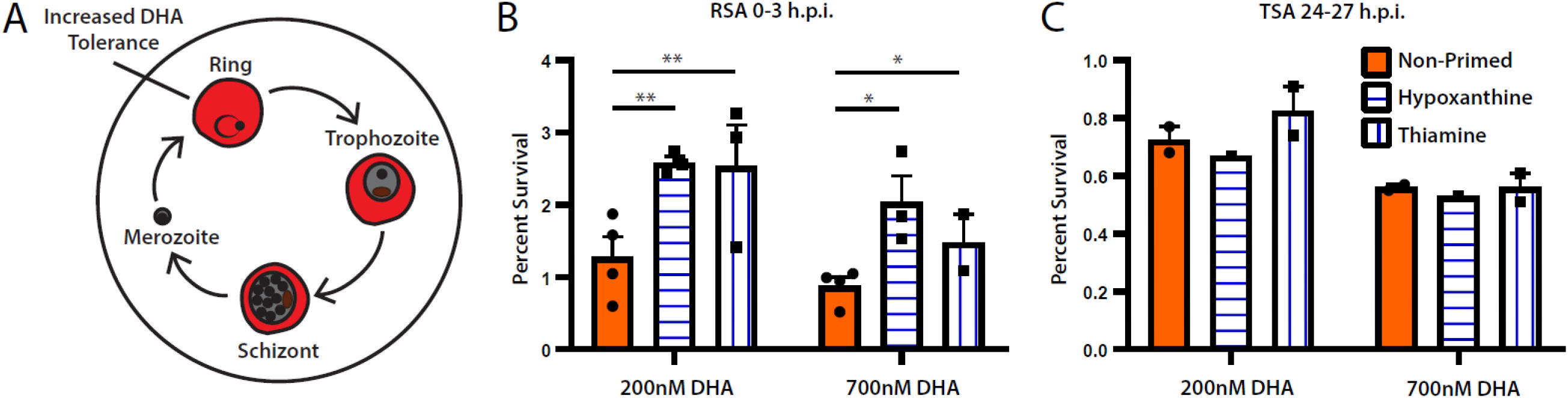
Increases in post-DHA survival are mediated by ring stage parasites. A) *P. falciparum* asexual blood stage replication cycle. DHA tolerance is highest in ring stage parasites 0-3h post-invasion^21^. B-C) Percent survival for early rings (B, 0-3h post-invasion, *N=*3-4) or trophozoites (C, 24-27h post-invasion, *N=*1-2) measured 66h after treatment with 200 or 700nM DHA for 6h. Bars represent S.E.M. Parasite line: *Dd2*. RSA p-values are as follows: Non-primed vs. low hypoxanthine: 200nM = 0.031, 700nM = 0.056. Non-primed vs. thiamine-free: 200nM = 0.038, 700nM = 0.073. TSA p-values are as follows. Non-primed vs. low hypoxanthine: 200nM = 0.407, 700nM = 0.930. Non-primed vs. thiamine free: 200nM = 0.796, 700nM = 0.999. P-values <0.1 are indicated with one asterisk (*). P-values <0.05 are indicated with two asterisks (**).

During metabolic priming experiments, the level of stress applied to the parasites impacted whether we observed superior recovery. For example, a >10% growth reduction over the course of low hypoxanthine metabolic priming was found to facilitate the phenotype (**Fig. 1C**), while failure to detect a growth phenotype (**Supplemental Fig. 3A**) or overly stressful conditions (**Supplemental Fig. 3B**) were detrimental to post-DHA recovery. These data support a biphasic dose response to metabolic priming characteristic of hormesis. However, this pattern is dependent on the limiting nutrient; in other conditions, growth during priming did not serve as a proxy for estimating dose response. For example, thiamine-free priming did not lead to growth reduction but still led to increased cumulative proliferation following DHA treatment (**Fig. 1D**). The differential effect of priming treatments on growth is likely due to parasite ability to scavenge or *de novo* synthesize thiamine, whereas *P. falciparum* are strictly purine auxotrophs.

**Figure 3.**
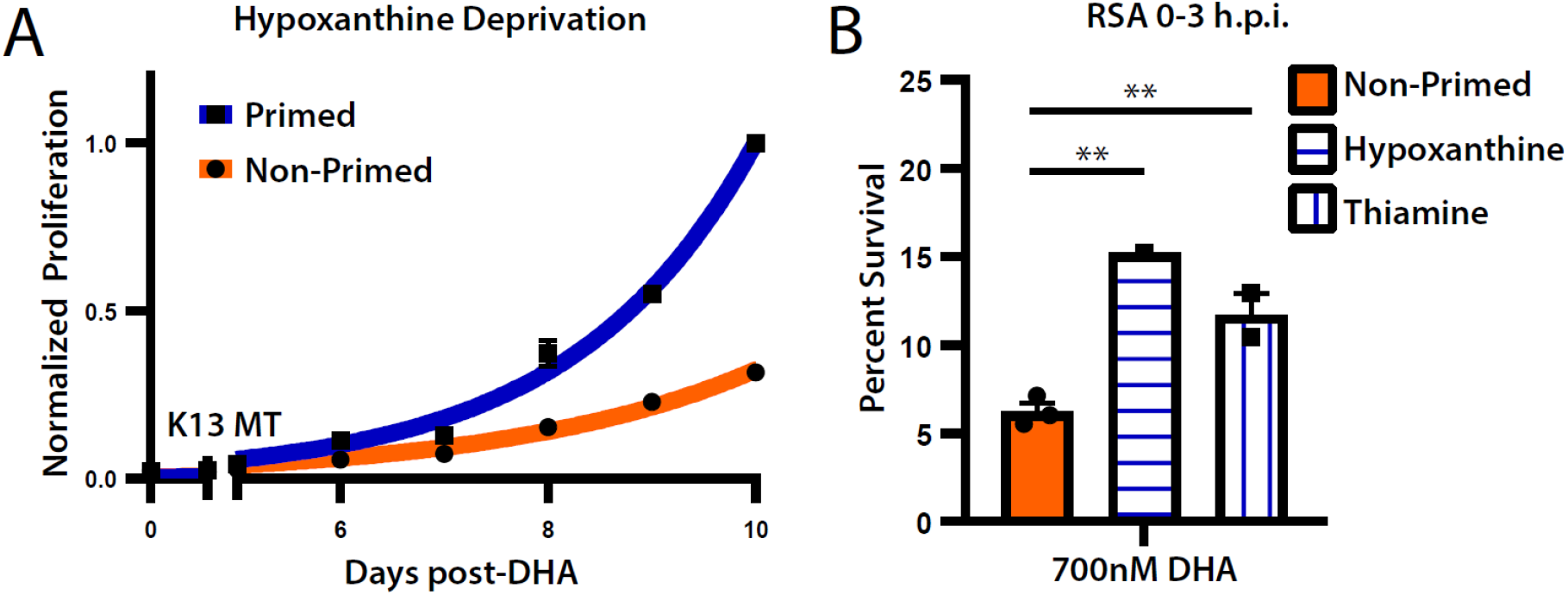
Metabolic priming effects are conserved in parasites carrying genetic DHA resistance. A) Post-DHA recovery of DHA tolerant parasite line in standard media and primed in low hypoxanthine or thiamine-free media. Bars represent S.E.M. of technical replicates within one representative experiment. B) Percent survival for early rings (0-3h post-invasion) measured 66h after treatment with 700nM DHA for 6h (*N*=1-3). Bars represent S.E.M. Parasite line: *MRA-1238*. RSA p-values are as follows: Non-primed vs. low hypoxanthine: 700nM = 0.012. Non-primed vs. thiamine free: 700nM = 0.026. P-values <0.1 are indicated with one asterisk (*). P-values <0.05 are indicated with two asterisks (**).

### Metabolic priming is stage-specific and facilitates increased DHA survival rates

DHA survival patterns are dependent on the parasite life cycle stage (with highest survival at 0-3hr post-invasion^21^, **Fig. 2A**). Therefore, we evaluated 1) the stage effects of metabolic priming and 2) the stage-specificity of the proliferation effect.

First, we compared parasite stage distribution in primed and non-primed populations to assess if subtle changes in lifecycle stage occur during priming conditions (i.e. stage effects). We did not detect any significant differences in the percentage of ring-stage parasites in non-primed compared to primed samples (Non-primed mean ± S.E.M.: 61.6% ± 2.1%, Primed: 61.3% ± 2.3%; *p*=0.67), (**Supplemental Table 1**). Therefore, an alteration of cell cycle progression is not contributing to differences in sensitivity to DHA treatment.

Second, we performed shorter-term DHA survival assays on highly synchronous parasite populations following priming to determine if changes in parasite recovery are stage-specific. To assess the behavior of early stage parasites (**Fig. 2A**), we performed “ring stage survival assays” (RSAs) on highly synchronous populations of 0-3h rings^21^. Metabolic priming prior to DHA treatment increased ring stage survival (**Fig. 2B**); we observed a mean 2.1-fold increase for both 200nM and 700nM DHA, which is the concentration used for our study (**Fig. 1C-D**) and the standard concentration for DHA RSAs, respectively. Notably, metabolic priming increased survival percentages to a level that is considered resistant (>1% for 700nM DHA for 6h^21^). Additionally, we tested later stage parasites (24-27h) in a modified RSA that assesses trophozoite survival (TSA^21^, **Fig. 2C**). We detected no significant increase in DHA survival following metabolic priming, indicating the benefit of metabolic priming is ablated in trophozoites (**Fig. 2D**). In addition to showing ring stage-specificity of the effect, these experiments also showed that higher cumulative proliferation in primed samples post-DHA is a result of an increased percentage of parasites surviving DHA treatment, as opposed to, increased proliferation per parasite.

### Metabolic priming increases survival in parasites with genetic DHA resistance

We sought to assess if the survival benefits of metabolic priming also apply to DHA resistant parasites (RSA >1%^21^). Using our long term growth assay, we found that metabolic-priming does improve post-drug recovery of a moderately DHA resistant parasite strain (**Fig. 3A**; MRA-1238, RSA: 6.2% survival, *kelch13* I543T mutant^22^). We confirmed this effect using RSAs; metabolic priming increases survival of a high DHA concentration (700mM) by a mean of 2.1-fold (**Fig. 3B**). This parasite line was not tested at the lower 200nM concentration due to their known resistance status; other DHA resistant lines (i.e. MRA 1240, RDA: 88.2% survival, *kelch 13* R539T) could not be assessed because of their high tolerance to 700nM DHA levels. These data in combination with data generated using DHA sensitive lines (**Figs. 1** and **2, Supplemental Table 1;** *Dd2* and *NF54*), indicate that increased DHA survival following metabolic priming triggers an intrinsic response that is not dependent on genetic background.

### Autophagy contributes to increased DHA survival

Phosphorylation of the translation initiation factor eIF2α occurs in *Plasmodium* in response to nutritional stress^23,24^. Additionally, increased levels of eIF2α phosphorylation are protective for DHA treatment^25^. In higher eukaryotes, phosphorylation of eIF2α can promote activation of the PI3K complex and increase transcription of autophagy-related genes^26,27^. In *Plasmodium*, increased PI3K activity mediates artemisinin resistance^28^. Further, polymorphisms in autophagy-related genes, such as ATG18 and ATG7, are associated with decreased artemisinin sensitivity in Southeast Asian parasite populations^29,30^. Recent evidence indicates the role of eIF2α phosphorylation in promoting autophagy is conserved in *Plasmodium*^31^. Thus, autophagy may be activated by metabolic priming in a manner associated with DHA survival.

To explore the role of autophagy in priming, we used a parasite strain with a conditional regulation of ATG8 expression; removal of the small molecule, anhydrotetracycline (aTc), leads to knockdown of ATG8, which is the essential for autophagosome formation and therefore essential for successful macroautophagy^32^. When we compared post-DHA recovery between parasites with low versus normal levels of ATG8 expression (**Supplemental Fig. 4 A-B**), we detected a 3.6-fold lower recovery benefit in parasites with low ATG8 expression (ATG8 On day 10 difference between low hypoxanthine and non-primed: 304%, ATG8 Off: 85%; **Supplemental Fig. 4C-D, Supplemental Table 1**). Along with translational repression, this result suggests that autophagy contributes to priming-induced pro-survival phenotype. However, the effect is difficult to fully ascertain using this approach, given that autophagy plays a role in the recovery of DHA treatment^29,31^; we confirmed this role by measuring the difference in post-drug recovery time between non-primed parasites with normal and low ATG8 expression. Parasites with normal ATG8 surpass 5% parasitemia ∼4 days earlier than parasites with low ATG8 expression following a 200nM DHA pulse (ATG8 On: 7.3d ± 0.7d (S.E.M.), ATG8 Off: 11.5d ± 0.5d, **Supplemental Fig. 4E**)

**Figure 4.**
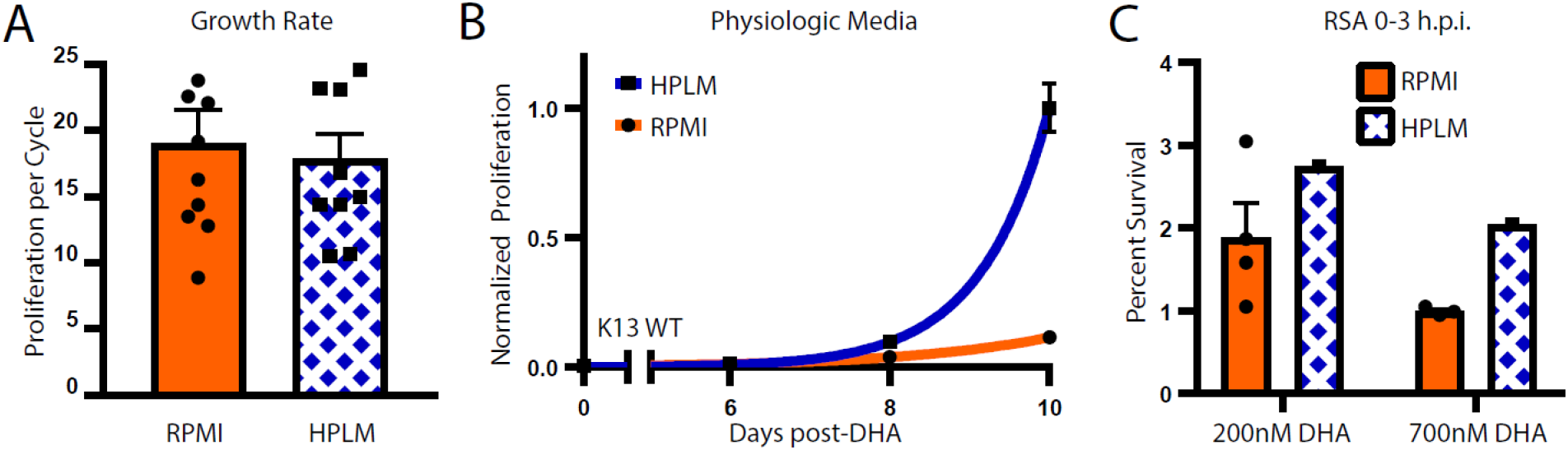
Physiologic media increases post-DHA recovery. A) Average growth rate of *Dd2* parasites propagated in RPMI-versus HPLM-based media. *N=*10 representing 5 replication cycles of two independent growth assays. B) Post-DHA recovery of *P. falciparum* propagated in RPMI- or HPLM-based media. Bars represent S.E.M. of technical replicates within one representative experiment. Results from independent assays are detailed in Supplemental Table 1 (*N=*2). C) Percent survival for early rings (0-3h post-invasion) measured 66h after treatment with 200 or 700nM DHA for 6h (*N*=1). Parasites were adapted to HPLM-based media at least 7 days prior to drug treatment and maintained in this media through the RSA reading. Bars represent S.E.M. Parasite line: *Dd2*. RPMI = Roswell Park Memorial Institute Media. HPLM = Human Plasma-Like Media.

### Physiological growth conditions promote increased DHA survival

Nutrient limitation is a physiologically relevant stress that parasites adapt to in the context of human infection. Our low hypoxanthine and thiamine-free metabolic priming conditions mimic nutrient stress from a single nutrient deficiency. While sole micronutrient deficiencies occur *in vivo*^33,34^, this is not the only relevant nutritional difference between *in vitro* cultures and *in vivo* infections. In general, levels of not just one, but many, nutrients are dramatically lower *in vivo* compared to the supra-physiological levels in standard RPMI-based media formulations^12-14^.

We hypothesized that the nutritional differences between RPMI and human plasma is sufficient to induce a metabolic priming-like effect similar to that seen with our established conditions. To test this, we utilized Human Plasma-Like Media (HPLM) as a base for *P. falciparum* culture^35^. Under routine propagation conditions (see *Methods*), parasites maintained in HPLM-based media grew equivalent to those in RPMI-based media (HPLM (mean ± S.E.M): 18 ± 1.9-fold per replication cycle, RPMI: and 19 ± 2.5-fold, **Fig. 4A**).

We adapted parasites to HPLM-based media for at least 7 days prior to treating with DHA and maintained parasites in the same media for 10 days of recovery. Propagation in HPLM-based media prior to DHA pulse led to superior post-drug recovery compared parasites in RPMI-based media (mean day 10 difference: 459%, *N=* 2, *p =* 0.0037, **Fig. 4B, Supplemental Table 1**). As with low hypoxanthine and thiamine-free priming conditions, growth in HPLM also increased DHA survival in RSAs (**Fig. 4C**, mean difference of 1.4-fold including both 200 and 700nM DHA samples). These data provide evidence that related pro-survival changes are occurring in response to both single nutrient deficiency and broader reductions of environmental nutrients.

*Metabolic priming shifts transcriptome toward artemisinin tolerance*

To more fully explore pathways that are altered in response to nutritional environment, we assessed gene expression in synchronous ring stage non-primed parasites and parasites primed with low hypoxanthine, thiamine-free, or HPLM conditions. Principal component analysis showed that each group had a distinct transcriptional profile, with low hypoxanthine primed parasites clustering separate from the other three groups on PC1 and the remaining groups separating on PC2 (**Fig. 5A**). This pattern parallels the number of significantly deferentially expressed (DE) genes detected in experimental groups compared to non-primed control samples with low hypoxanthine primed parasites yielding the largest number of differences (**Fig. 5B**).

**Figure 5.**
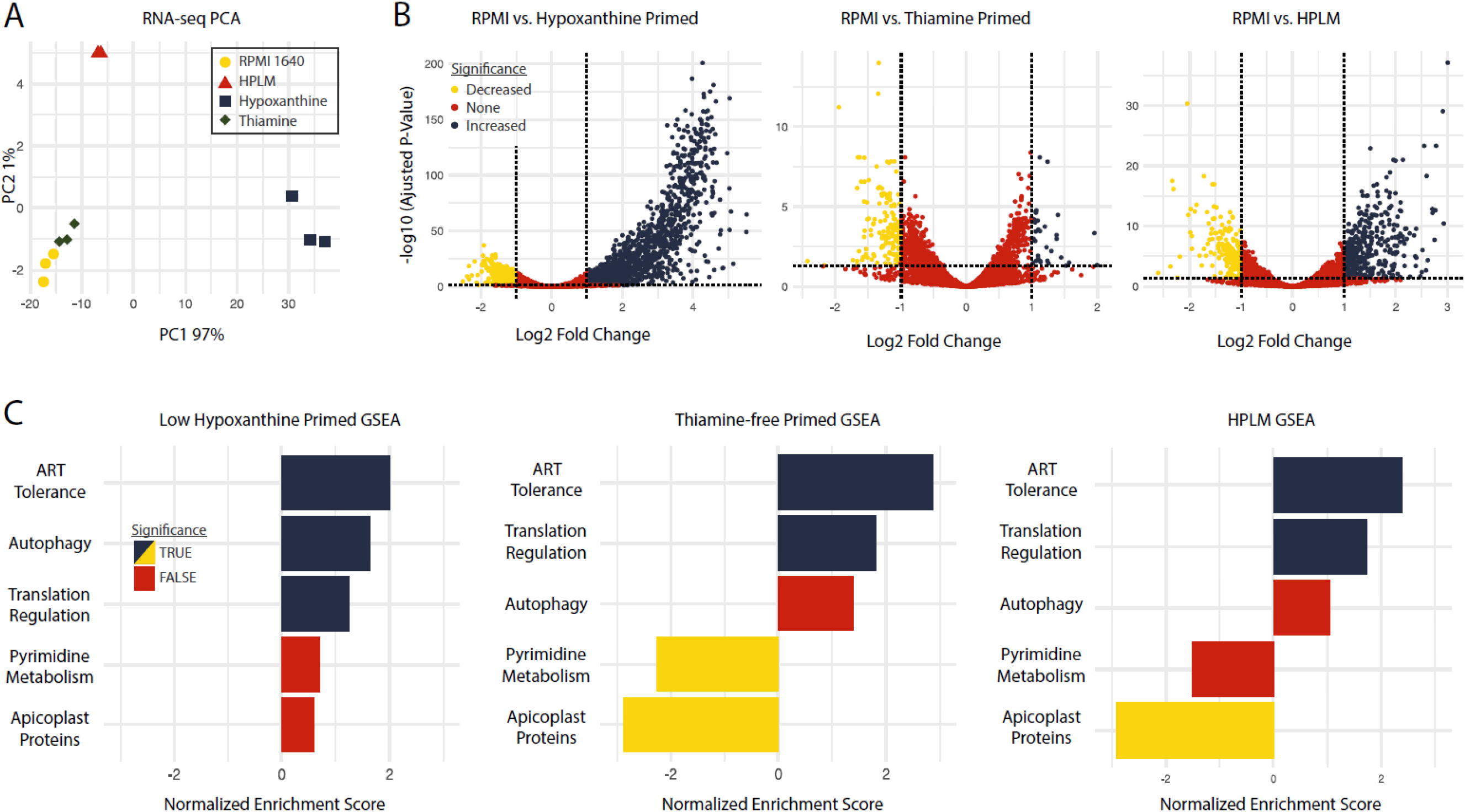
Metabolic priming induces artemisinin tolerance-like transcriptional changes. A) PCA plot of gene expression from each condition. B) Volcano plots of differentially expressed genes for each condition. Log2 fold change thresholds set at - 1 and +1. Adjusted p-value set to ≤0.05. C) GSEA results for selected pathways. TRUE and FALSE = adjusted p-value of <0.05 and >0.05, respectively. RPMI = Roswell Park Memorial Institute Media. HPLM = Human Plasma-Like Media. GSEA = Gene Set Enrichment Analysis. PCA = Principal Component Analysis.

To identify pro-survival pathways, we performed Gene Set Enrichment Analysis (GSEA) across priming conditions (**Fig. 5C**). Some metabolic pathways not previously implicated in stress responses, but important for parasite biology, were altered in a condition-dependent manner.

For example, pyrimidine biosynthesis and apicoplast metabolism were significantly down-regulated in in thiamine-free and HPLM groups, respectively (**Fig. 5C**). Despite the impact of ATG8 knockdown on our phenotype (**Supplemental Fig. 4C-D**), GSEA indicated autophagy was significantly upregulated in low hypoxanthine primed samples but not across all groups (**Fig. 5C**). Thus, induction of autophagy may be condition-specific and is unlikely to be a primary driver of the observed phenotype.

Overall, GSEA showed that 25 of 463 total pathways were significantly enriched across all groups (**Supplemental Table 2**, including pathways heavy in exported proteins (*e*.*g*. protein-protein interactions between *P. falciparum* and the red blood cell). Upregulation of exported proteins has been previously observed in both the artemisinin resistance transcriptome and as a response to nutrient limitation^36-38^. Translational regulation was also significantly upregulated in all three conditions in concordance with previous knowledge of the eIF2α-mediated response under nutrient stress (**Fig. 5C, Supplemental Table 2**). This pathway includes genes with roles in processes known to impact DHA survival including the unfolded protein response (UPR) and proteasomal degradation (**Supplemental Table 2**).

**Table 2.** Genes of interest curated from differentially expressed gene lists. Asterisk indicates genes found to be basally altered in artemisinin mutants (from **Supplemental Table 3**).

| Gene ID | Product Description |
| --- | --- |
| *Upregulated* |  |
| PF3D7_0214600* | serine/threonine protein kinase STK2, putative |
| PF3D7_0402200 | surface-associated interspersed protein 4.1 (SURFIN 4.1), pseudogene |
| PF3D7_0418600 | regulator of chromosome condensation, putative |
| PF3D7_0613900 | myosin E, putative |
| PF3D7_0618000 | conserved Plasmodium membrane protein, unknown function |
| PF3D7_0707300 | rhoptry-associated membrane antigen |
| PF3D7_0724900 | kinesin-20, putative |
| PF3D7_0817600* | conserved protein, unknown function |
| PF3D7_0822900* | PhIL1-interacting candidate PIC2 |
| PF3D7_0903600 | conserved protein, unknown function |
| PF3D7_0911100 | START domain-containing protein, putative |
| PF3D7_0919900 | regulator of chromosome condensation-PP1-interacting protein |
| PF3D7_1026600 | conserved Plasmodium protein, unknown function |
| PF3D7_1035300* | glutamate-rich protein GLURP |
| PF3D7_1126700 | conserved Plasmodium protein, unknown function |
| PF3D7_1143100 | AP2 domain transcription factor AP2-O |
| PF3D7_1145200* | serine/threonine protein kinase, putative |
| PF3D7_1206300* | conserved Plasmodium protein, unknown function |
| PF3D7_1223100* | cAMP-dependent protein kinase regulatory subunit |
| PF3D7_1251200* | coronin |
| PF3D7_1252400 | reticulocyte binding protein homologue 3, pseudogene |
| PF3D7_1327300 | conserved Plasmodium protein, unknown function |
| PF3D7_1335400 | reticulocyte binding protein 2 homologue a |
| PF3D7_1356800 | serine/threonine protein kinase ARK3, putative |
| PF3D7_1371600 | erythrocyte binding like protein 1, pseudogene |
| PF3D7_1423300 | serine/threonine protein phosphatase 7 |
| *Downregulated* |  |
| PF3D7_0308000 | DNA polymerase delta small subunit, putative |
| PF3D7_0510100 | KH domain-containing protein, putative |
| PF3D7_0512800 | conserved Plasmodium protein, unknown function |
| *PF3D7_0711000* | AAA family ATPase, CDC48 subfamily |
| *PF3D7_0715900* | cation diffusion facilitator family protein, putative |
| *PF3D7_0801100* | 28S ribosomal RNA |
| *PF3D7_0802200* | 1-cys peroxiredoxin |
| *PF3D7_0923800* | thioredoxin reductase |
| *PF3D7_1130900* | conserved Plasmodium protein, unknown function |
| *PF3D7_1239200* | AP2 domain transcription factor, putative |
| *PF3D7_1303800* | conserved Plasmodium protein, unknown function |
| *PF3D7_1322300* | BRCT domain-containing protein, putative |
| *PF3D7_1330600* | elongation factor Tu, putative |
| *PF3D7_1334500* | MSP7-like protein |
| *PF3D7_1359600* | conserved Plasmodium protein, unknown function |
| *PF3D7_1371800* | Plasmodium exported protein, unknown function |
| *PF3D7_1419200* | thioredoxin-like protein, putative |
| *PF3D7_1440200* | stromal-processing peptidase, putative |
| *PF3D7_1443900* | heat shock protein 90, putative |
| *PF3D7_MIT04100* | large subunit ribosomal RNA fragment D |

Due to the large number of DE genes following hypoxanthine priming, and relatively small number in thiamine-free samples, it was necessary to limit the complexity of our results to further assess individual transcripts contributing to our phenotype. We specifically focused on the top 15 up- and down-regulated genes in each group as determined by lowest adjusted p-values. This yielded 37 up- and 42 down-regulated unique genes across the three treatment conditions as there was some overlap between groups. From this list of genes, we identified those that had an adjusted p-value of ≤0.05 in all groups as a curated list of 26 up- and 20 down-regulated genes of interest (**Table 2**). This curated list contained many genes implicated in artemisinin genetic tolerance, such as *coronin* (PF3D7_1251200), a gene known to be sufficient to induce artemisinin tolerance when mutated^39^. When compared to a list of 151 genes known to be basally altered (without active DHA treatment; **Supplemental Table 3**) in genetically artemisinin tolerant parasites^39-43^, 8 of our curated genes were represented (**Table 2**). Further, changes in this curated set of 151 genes (designated as “ART Tolerance” in **Fig. 5C**) were highly enriched by GSEA in all groups (in the top 1% of 463 pathways by adjusted p-value). These data indicate that nutrient limitation remodels metabolism in parasites to induce an artemisinin tolerance-like state.

## Discussion

In the current study, we have demonstrated that physiologically relevant nutrient limitation stimulates intrinsic pro-survival pathways in *P. falciparum* (**Figs. 1-4**). Our analyses indicate genes within these altered pathways directly contribute to genetic artemisinin tolerance (**Fig. 5C**), thus we speculate that nutrient limitation offers a preamble for resistance evolution. Our findings connect to other findings on cellular responses to both nutrient limitation and tolerance (**Fig. 6**) and provide a framework for how the nutritional environment impacts pathogen homeostasis. We explore these connections and their relevance below.

**Figure 6.**
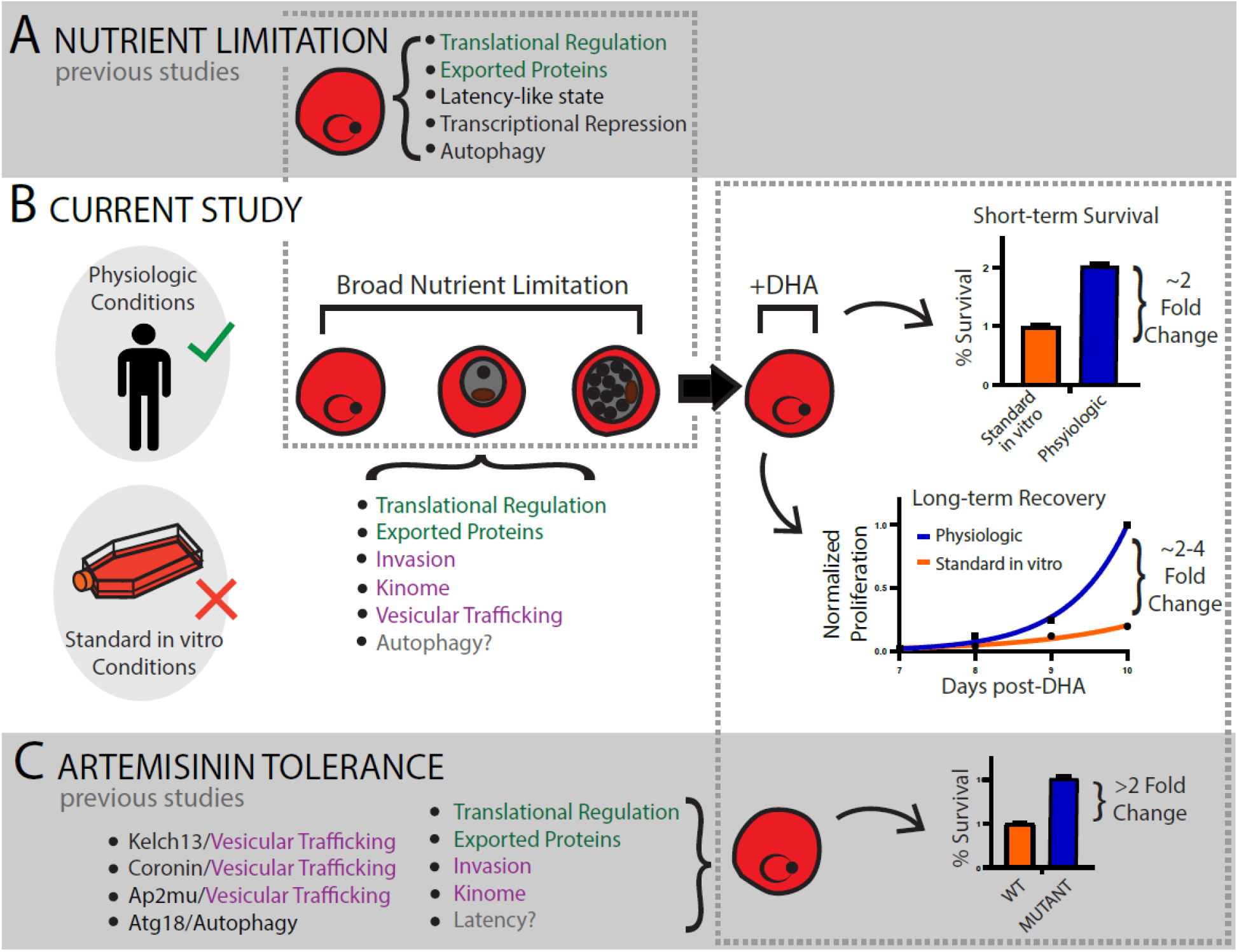
Integration of information related to nutrient deprivation and artemisinin tolerance. **A**) Previous studies identified that nutrient limitation has metabolic impacts on *Plasmodium* parasites. Translational regulation: eif2α^24,31^; Exported proteins^37^; Latency-like state^23^; Transcriptional Repression^44^; Autophagy: ATG5/8^58^. **B**) The current studies have identified a direct link between nutrient limitation and parasite survival following dihydroartemisinin (DHA) treatment. These conditions are more relevant in the human host than in standard *in vitro* conditions. Transcriptional changes induced by nutrient limitation mimic those that contribute to artemisinin tolerance. Translational regulation: Fig. 5C, Supplemental Table 2, “Transl_Regul” (285 genes); Exported proteins: Supplemental Table 2 “Meroz_RBClig” (19 genes) and “DomSurfaceProt” (14 genes); Invasion: Table 2 (i.e. GLURP, Phil1), Supplemental Table 2 “Erythrocyteinvasionpath” (8 genes), “Merozoiteproteins” (89 genes); Kinome: Table 2 (i.e. STK2, STK, CDPK, ARK3), Supplemental Table 2 “Phospho_mero” (482 genes); Vesicular trafficking: Table 2 (i.e. coronin); Autophagy: Supplemental Fig. 4 (ATG8), Fig. 5C (only Hypoxanthine priming), grey text-full contribution to phenotype remains unclear due to the contribution of autophagy to DHA recovery^29,31^, Supplemental Fig. 4. **C**) Previous studies identified pathways associated with artemisinin tolerance. Vesicular trafficking: kelch13^28^, coronon^39^, Ap2μ^59^; Autophagy: ATG18^29,31^; Translational regulation: unfolded protein response^60^, proteasomal degradation (UBP1^59^); Exported proteins^36,38^; Invasion: GLURP and Phil1^40^; Kinome: STK2, STK, CDPK^40^, Latency: grey text-unclear if distinct from nutrient-dependent latency or directly contributes to tolerance^25,61^.

### Limiting nutrient levels for malaria parasites in vitro

The supra-physiological concentrations of nutrients found in common media formulations, such as RPMI-1640, can potentiate genetic, transcriptional, and metabolic changes in parasite biology (reviewed in ^12,14^). Here, we observed superior parasite recovery following limitation of single nutrients in *in vitro* growth media; hypoxanthine and thiamine were chosen in part due to contrasting metabolic attributes outlined in the *Results* section, but also due to potential relevance to conditions *P. falciparum* may experience *in vivo* (see further discussion below). This approach is similar to other studies that induced nutrient stress by reduction of a single amino acid^23,44^ or select metabolite classes^24,31,45^. We also observed superior recovery when parasites were grown in HPLM, a physiological media formulation that has reduced concentrations of multiple nutrients^13^. To our knowledge, this is the first use of HPLM as a media base for the propagation of *P. falciparum*; to induce starvation in previous studies, nutrient-rich RPMI-based media is diluted^45^, producing a blanket reduction in nutrient levels. Regardless of the approach, the *P. falciparum* response to artemisinin treatment, when not protected by this supra-physiological glut of nutrients, had not been previously evaluated. This study begins to connect disparate observations to improve our understanding of how the environment impacts cellular responses.

### Molecular connections between nutrient limitation and DHA survival

We demonstrate that metabolic priming triggers broad alterations of translational regulation (**Fig. 5C** and **6B**), including genes associated with the UPR and proteasomal degradation. Previous studies in *Plasmodium* connected nutrient limitation with translational regulation (via eIF2α phosphorylation^23,24^) and separately, eIF2α phosphorylation with survival of DHA treatment ^25^ (**Fig. 6A** and **C**). Our study establishes a direct link between nutrient limitation and DHA survival (**Fig. 6B**). Additionally, our data support a role for autophagy in priming-induced DHA survival (**Supplementary Fig. 4C-D** and **Fig. 5C**). Given that the activation of autophagy in *P. falciparum* and higher eukaryotes is partially dependent on translational regulation by eIF2α phosphorylation^31,46^, the connection between nutrient limitation, eIF2a-autophagy, and DHA survival is growing stronger. However, the extent to which eIF2α phosphorylation-dependent latency is mediating this phenotype remains to be explored.

A surprising outcome of our study was the discovery that nutrient limitation triggered many responses that mimic an artemisinin tolerance-like transcriptional state (**Fig. 6B** and **C**). This adaptive response, which was stimulated by all priming conditions (**Fig. 5**), triggered a change in expression for genes that had previously been linked to genetic artemisinin tolerance (**Table 2**). Future investigations should determine the longevity of priming-induced pro-survival signals and whether this adaptation facilitates development of genetic tolerance. This phenomenon has been observed in the development of antibiotic resistant bacteria^47^ and recently proposed for clinical *Plasmodium* artemisinin resistance^38^. Of note, more than half of the top transcriptional markers for resistance identified by Zhu, *et al*. were significantly altered (by adjusted p-value) in the same direction in at least one of our priming conditions (32/58 genes).

### Translation from in vitro to in vivo results

Analyses of how the *in vitro* environment affects pathogens outside of their native environment remains limited. Alterations of metabolism that influence pathogen drug sensitivity can lead to discordance between *in vitro* results and *in vivo* responses. Our data emphasizes the importance of the nutritional environment, whether standard *in vitro* culture conditions or more *in vivo*-like environments, on malaria parasite survival (**Fig. 6B**). Moving forward, the use of physiological HPLM-based media may be appropriate for select assessments where clinical translation is paramount. Future studies should also investigate if the nutritional environment is relevant to other antimalarial drugs and how pro-survival pathways can be targeted to enhance their killing potential.

### Relevance for the in vivo host environment

Increasing resistance to artemisinin and its partner drugs necessitates development of new antimalarial drugs. Nucleotide metabolism is a potential future drug target in *P. falciparum*; antimalarials are currently being developed to inhibit both purine scavenging (including hypoxanthine) and pyrimidine biosynthesis^48-51^. As some of these candidates move closer to use in the clinic, it is imperative to understand how parasites respond to the stress of nucleotide deprivation. Similarly, thiamine limitation has physiological relevance to *P. falciparum* infection as clinical thiamine deficiency is common in malaria endemic regions, such as Southeast Asia^33,34^, and evidence suggests malaria infection exacerbates this deficiency^52^. Additionally, metabolic network reconstructions imply that drug resistance status can alter metabolism, making parasites more reliant on thiamine import for survival^53^. This metabolic shift is corroborated by increased expression in artemisinin tolerant isolates of thiamine pyrophosphokinase, a gene necessary for conversion of imported thiamine to the active thiamine pyrophosphate form^54^.

Combining artemisinin with partner drugs of distinct mechanisms of action has contributed to the effectiveness of antimalarial drugs over recent decades. Such combination therapies minimize the development of tolerance or resistance. Yet, the potential for partner drugs that target metabolism (as described above) to induce an artemisinin tolerant-like transcriptional state poses an unexpected challenge that should be considered in the development of new antimalarial combinations.

Combating antimalarial drug resistance is especially critical as the spread of genetic artemisinin tolerance threatens to undo progress in reducing malaria mortality^55^. However, the challenges of fighting drug resistance are not unique to malaria. The interplay between nutrient limitation, pro-survival pathways, and sensitivity to drug treatment (**Fig. 6**) has important implications for the evolution of drug resistance across diverse pathogens. How microbes adjust their metabolism in ways that impact survival of current and future drugs is an important avenue of research to prevent and slow the spread of drug resistant pathogens.

## Materials and Methods

### Parasites and Growth

*Plasmodium falciparum* lines MRA 150 (Dd2), MRA 1000 (NF54), and MRA 1238 were obtained from the Malaria Research and Reference Reagent Resource Center (MR4, BEI Resources). The ATG8 conditional TetR-Dozi knockdown line was a generous gift from Ellen Yeh (Stanford University, Stanford, CA). *Plasmodium* cultures were maintained in A+ human erythrocytes (Valley Biomedical, Winchester, VA) at 3% hematocrit in RPMI 1640 HEPES (Sigma Aldrich, St Louis, MO) supplemented with 0.5% Albumax II Lipid-Rich BSA (Sigma Aldrich, St Louis, MO) and 50 mg/L hypoxanthine (Thermo Fisher Scientific, Waltham, MA). This formulation is referred to as “standard media” throughout the manuscript. ATG8 TetR-DOZI parasites were maintained with 0.5μM anhydrotetracycline (Sigma Aldrich, St Louis, MO) or supplemented with isopentyl pyrophosphate (Isoprenoids LC, Tampa, FL) as previously described^32^. Cultures were grown at 37°C and individually flushed with 5% oxygen, 5% carbon dioxide, 90% nitrogen gas. Dilution of cultures with uninfected erythrocytes and changing of culture medium was performed every other day. Parasitemia was determined by flow cytometry using SYBR Green I staining. Cultures were confirmed negative for mycoplasma approximately monthly using a LookOut Mycoplasma PCR detection kit (Sigma Aldrich, St Louis, MO).

### Metabolic Priming

Low hypoxanthine media was made using RPMI 1640 HEPES base with reduced exogenous hypoxanthine addition to a final concentration of 0.5μM. Thiamine-free media was custom ordered to be identical to RPMI 1640 HEPES without the addition of thiamine hydrochloride. Hypoxanthine was added to thiamine-free media to 50 mg/L. Human Plasma-Like Media (HPLM, Thermo Fisher Scientific, Waltham, MA) was used without added HEPES or exogenous hypoxanthine. All media formulations were supplemented with 0.5% Albumax II Lipid-Rich BSA.

Uninfected erythrocytes were pre-incubated prior to use in priming experiments in the appropriate low nutrient medium for 48h at 37°C, 3-6% hematocrit, and 5% oxygen, 5% carbon dioxide, 90% nitrogen gas. Pre-incubated uninfected erythrocytes were seeded with infected erythrocyte culture to a starting parasitemia of <0.5% and resuspended in control medium for non-primed samples or low nutrient medium for primed samples. Cultures were allowed to incubate for the prescribed period of time dependent on the nutrient deprived (**Table 1**). For 72h incubations, media was refreshed at 48h.

Aliquots were taken for flow cytometry measurement with SYBR Green I and MitoProbe DiIC1(5) kit (both Thermo Fisher Scientific, Waltham, MA). Primed samples were compared to non-primed counterparts for reduction in growth, mitochondrial membrane potential (MMP), and percentage of ring stage parasites. Priming was considered successful when the following conditions were met for low hypoxanthine primed samples compared to non-primed: 1) an approx. >10% growth decrease, 2) little to no decrease in MMP (approx. <10% drop), and 3) ring stage percentages within approx. 10% of non-primed. Priming in thiamine-free and HPLM conditions was considered successful if both conditions “2” and “3” were met. Following successful priming, both primed and non-primed samples were treated with either 200nM dihydroartemisinin (DHA, Sigma Aldrich, St Louis, MO) or DMSO for 6h. To remove drug, cultures were washed three times in media and all samples were resuspended in normal, full-nutrient RPMI + Albumax media and returned to the incubator. Cultures were measured every other day with SYBR Green I and MitoProbe for 10-14 days post drug treatment.

Biological replicates were performed in technical duplicate. For statistical analysis, growth data 10 days post-DHA was log transformed and tested for normality by Shapiro-Wilks test prior to running a repeated measures two-way ANOVA followed by Sidak’s multiple comparisons test. RSA and TSA data was tested by ordinary two-way ANOVA followed by Dunnett’s multiple comparisons test. Analysis and visualizations were made using GraphPad Prism version 7.04.

### Ring and Trophozoite Stage Survival Assays

Survival assays were performed as previously described with minor modifications^56^. Briefly, all metabolically primed samples were suspended in the relevant media at 36h, 72h, or ≥7d prior to drug treatment for thiamine-, hypoxanthine-, and HPLM-primed conditions, respectively. Approximately 30h prior to 0-3h ring generation, 35mL cultures were synchronized with 5% D-sorbitol and allowed to progress until the culture was predominantly schizonts. Schizonts were isolated by layering 4 mL of culture over 4 mL of 75% Percoll, centrifugation, and collection of the intermediate band. Isolated schizonts were washed then added to uninfected erythrocytes in the appropriate media formulation. Exactly three hours later, a rapid D-sorbitol synchronization was performed (10m at 37 °C followed by 5s vortex) to remove any uninvaded late-stage parasites. Parasites for TSAs were returned to the incubator for exactly 24h before drug treatment, whereas parasites for RSAs were immediately pulsed with either DMSO or DHA (200nM or 700nM) for 6h. Following drug treatment, all cells were washed multiple times and returned to RPMI + ALB media for 66h prior to assessment of survival by flow cytometry measurement with SYBR Green I and MitoProbe DiIC1(5) kit.

### Immunoblotting

Anti-ATG8 antibody was a gift from Ellen Yeh (Stanford University, Stanford, CA). Anti-ATG8 was used in conjunction goat Anti-guinea pig IgG H&L (Alexa Fluor® 488) (Abcam, Waltham, MA). Bound antibodies were detected on a Bio-Rad Imager and quantified using Image Lab Software (Bio-Rad, Hercules, CA).

### RNAseq

Ring stage samples were generated by synchronization with 5% sorbitol 88h and 44h prior to harvest to accommodate the ∼44h erythrocytic cycle duration of Dd2^57^. Primed sample groups were grown as described above in their designated media for 72h, 36h, and 7d prior to extraction for hypoxanthine, thiamine, and HPLM groups, respectively. Starting 24h prior to RNA extraction, blood smears were assessed every 3 hours until reinvasion was observed to approximate the age of rings in final samples. In all RNAseq samples, reinvasion was first observed 18h prior to harvest. At sample collection, erythrocytes were lysed in 0.15% saponin before RNA was extracted from parasite pellets using a Direct-zol RNA Miniprep Kit (Zymo Research) according to the manufacturer’s instructions. An aliquot of each sample was taken for assessment with a 2100 Bioanalyzer using the RNA 6000 Pico assay (Agilent Technologies, USA) before the remainder of sample was snap frozen in liquid nitrogen and stored at -80C.

Samples were shipped to Genewiz, Inc. for standard RNAseq with polyA selection. Raw reads obtained from Genewiz, Inc. were checked for overall quality and trimmed to remove adapters and low-quality bases from 3’ ends (with parameter -q 25) using TRIMGALORE (v. 0.6.7). STAR (v. 2.7.9a) was used to align trimmed reads with parameters – genomeSAindexNbases 11 and – alignIntronMax 500. Featurecounts (v. 2.0.1) was used for gene counting with parameter -t ‘exon’. Differential expression analysis was done using DESeq2 (v. 1.34.0). Genes were considered significantly differentially expressed if the following criteria were met 1) Benjamini-Hochberg adjusted p-value of ≤0.05 and 2) log2-fold change of ≥1.0 or ≤ -1.0. Gene set enrichment analysis (GSEA) was performed using fgsea (v. 1.20.0). 3D7 genome sequence for alignment and annotation, gene product descriptions, and metabolic pathway information were pulled from PlasmoDB (Release 55). Visualizations and analyses were generated in R version 4.1.2 using Tidyverse (v. 1.3.1) packages.

