## Supplemental for "Nutrient limitation mimics artemisinin tolerance in malaria"

**Supplemental Information**

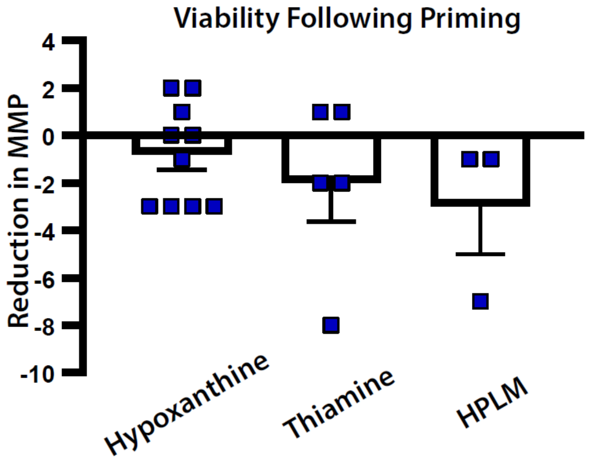

**Supplemental Figure 1. Metabolic priming does not drastically impact parasite viability.** Successful low nutrient metabolic priming leads to very small decreases in viability compared to standard media, non-primed controls. The percentage of parasites (SYBR Green I cells) also positive for MitoProbe DiIC1(5) staining (an indicator of mitochondrial membrane potential; MMP) was used as a proxy for determining viability. *N*=3-10 per condition. Bars represent S.E.M.

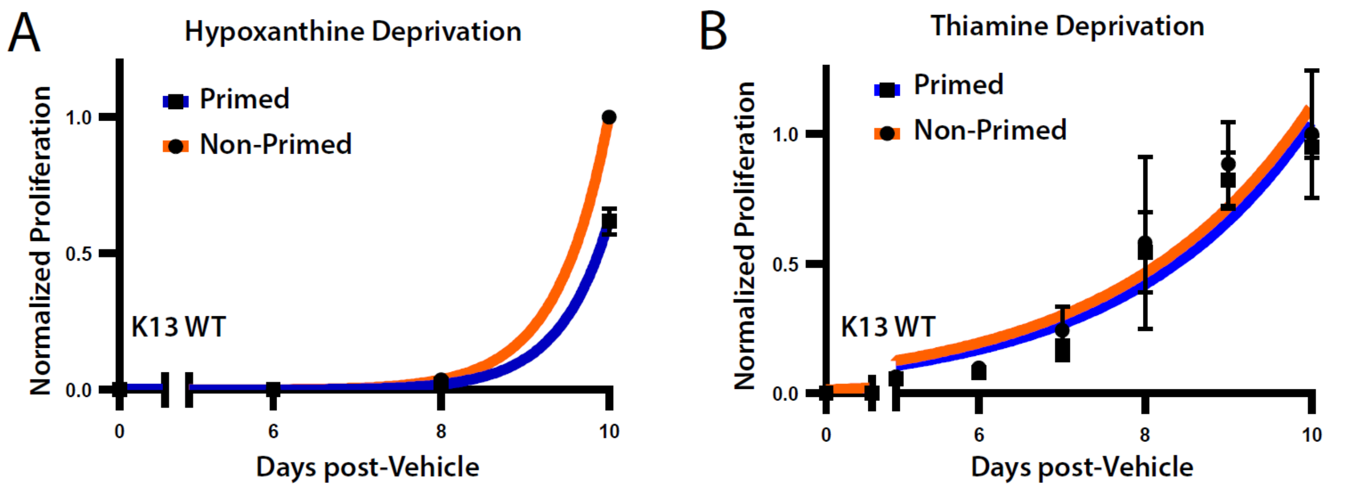

**Supplemental Figure 2. Metabolic priming does not impact post-DHA recovery without drug pulse.** A-B) Growth in standard media following metabolic priming under A) low hypoxanthine or B) thiamine-free conditions (Table 1) without a DHA drug pulse (vehicle is DMSO). Bars represent S.E.M. of technical replicates within one representative experiment. Results from independent assays are detailed in Supplemental Table 1 (*N*=2 per condition).

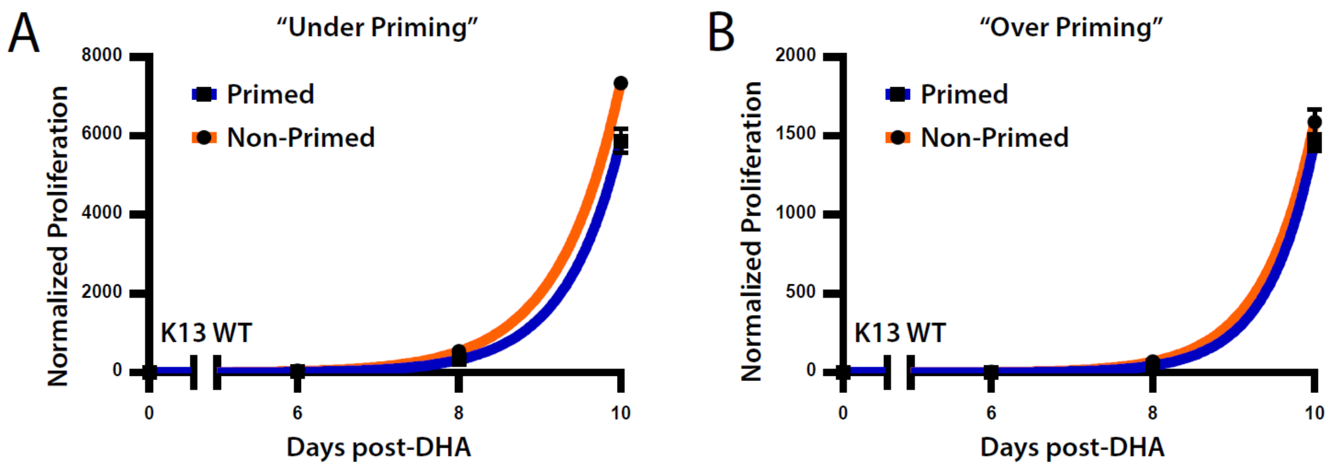

**Supplemental Figure 3. Increased post-DHA recovery requires a specific range of stress during metabolic priming.** A-B) Post-DHA recovery in standard media following low hypoxanthine metabolic priming where priming led to A) less than 10% growth reduction or B) greater than ~60% growth reduction and >10% reduction in MMP compared to non-primed controls. Bars represent S.E.M. of technical replicates within one independent experiment.

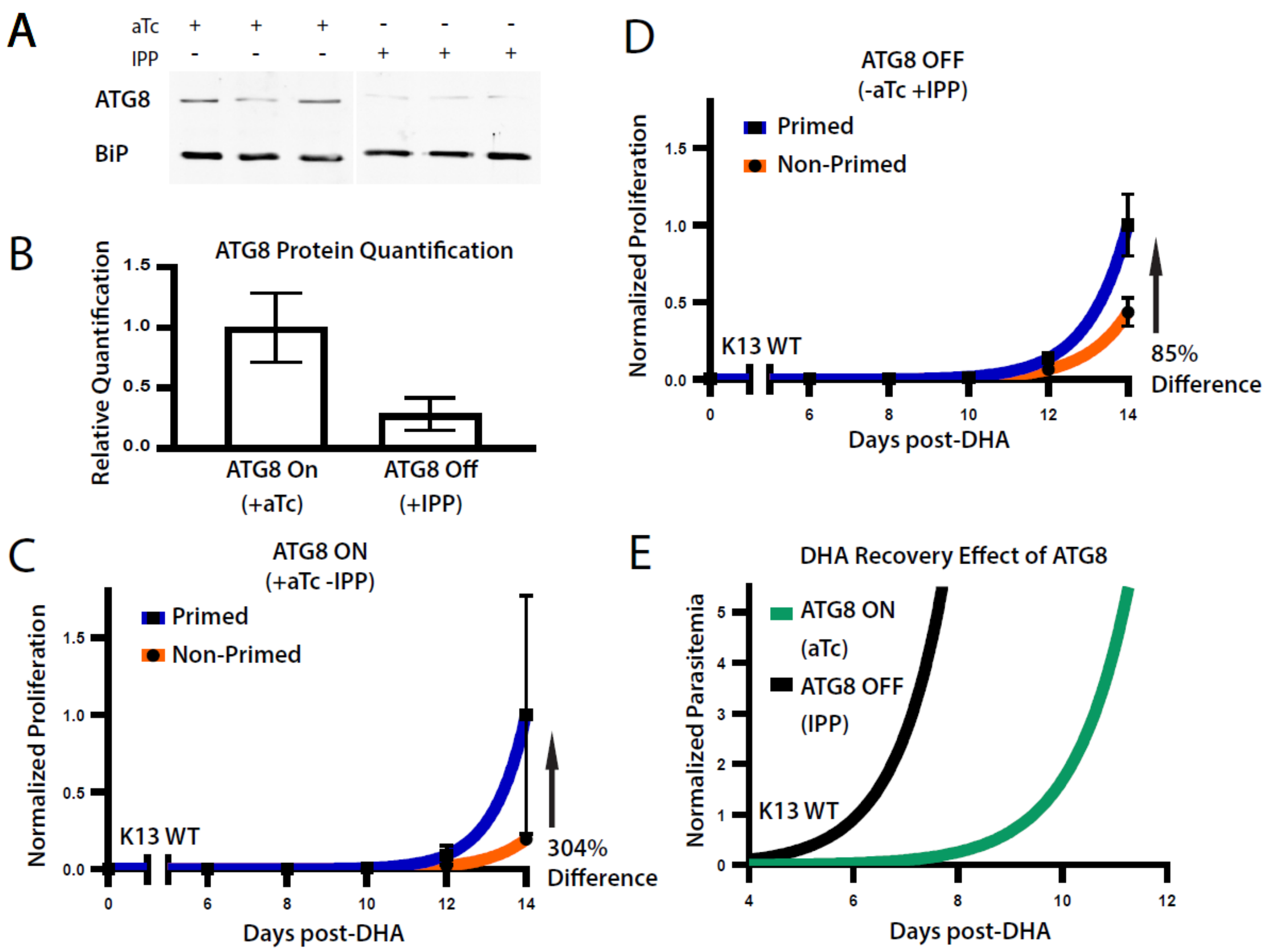

**Supplemental Figure 4. Autophagy plays a role in priming induced DHA tolerance.** A-B) Effect of knockdown on ATG8 protein levels in ATG8 TetR-Dozi parasites. Bars represent S.E.M. (*N*=3). C-D) Post-DHA recovery in a paired experiment of hypoxanthine primed ATG8 TetR-Dozi parasites with ATG8 expression on (C) or off (D). Bars represent S.E.M. of technical replicates within one independent experiment. E) Growth recovery of non-primed parasites differing in ATG8 expression status (*N*=2). aTc = anhydrotetracycline; small molecule that allows induction of ATG8 expression. IPP = isoprenoids; necessary metabolite for *P. falciparum* survival in the absence of ATG8.

**Supplemental Table 1.** Data for all recovery post-DHA or post-vehicle control experiments. Asterisk indicates experiment did not meet criteria for successful priming (**Supplementary Fig. 3**)

| LINE | PRIMING STRESS | DRUG STRESS | D10% DIFFERENCE | NON-PRIMED %RINGS | PRIMED %RINGS | PRIMING GROWTH REDUCTION | PRIMING MMP CHANGE | BLOOD AGE AT START (DAYS) |
|---|---|---|---|---|---|---|---|---|
| **DD2** | Hypoxanthine | DHA | 200 | 53.5 | 59 | 30 | n.d. | 14 |
| **DD2** | Hypoxanthine | DHA | 99 | 58.5 | 53.5 | 42 | -3 | 24 |
| **DD2** | Thiamine Free | DHA | 104 | 72.5 | 68.5 | -69 | -18 | 14 |
| **DD2** | Thiamine Free | DHA | 137 | 56.5 | 57 | -1 | 1 | 18 |
| **DD2** | HPLM | DHA | 676 | 73.5 | 74.5 | -4 | -7 | 28 |
| **DD2** | HPLM | DHA | 151 | 76 | 78 | 0 | -1 | 16 |
| **NF54** | Hypoxanthine | DHA | 20 | 47.5 | 45.5 | 14 | -3 | 27 |
| **NF54 (ATG8 ON)** | Hypoxanthine | DHA | 304 | 61 | 54 | 60 | 2 | 24 |
| **NF54 (ATG8 OFF)** | Hypoxanthine | DHA | 85 | 67 | 61 | 36 | 2 | 24 |
| **MRA1238 (K13 MT)** | Hypoxanthine | DHA | 142 | 66.5 | 63 | 36 | n.d. | 9 |
| **NF54** | Thiamine Free | DHA | 195 | 68.5 | 67.5 | 10 | 1 | 18 |
| **NF54** | Thiamine Free | DHA | 257 | 44.5 | 47 | 17 | -2 | 18 |
| **NF54** | Hypoxanthine | No Drug | -30 | 47.5 | 45.5 | 14 | -3 | 27 |
| **NF54** | Hypoxanthine | No Drug | -27 | 68.5 | 75 | 45 | 0 | 9 |
| **DD2** | Hypoxnathine | No Drug | -38 | 60 | 56 | 24 | -1 | 12 |
| **NF54** | Hypoxnathine | No Drug | -9 | 60.5 | 58.5 | 37 | -3 | 27 |
| **DD2** | Thiamine Free | No Drug | -5 | 48.5 | 44.5 | 7 | -2 | 16 |
| **DD2** | Thiamine Free | No Drug | 29 | 63 | 67 | -2 | 3.5 | 14 |
| **DD2** | HPLM | No Drug | 111 | 76 | 76 | -14 | n.d. | 21 |
| **DD2** | HPLM | No Drug | | 76 | 78 | 0 | -1 | 16 |
| ***NF54 (ATG8 ON)** | Hypoxanthine | DHA | -20 | 72 | 68 | 9 | 0 | 16 |
| ***D6** | Hypoxanthine | DHA | -8 | 62 | 32 | 66 | 1 | 20 |

**Supplemental Table 2.** Gene Set Enrichment Analysis Results (in separate excel file)

**Supplemental Table 3.** Studies measuring basal differences between artemisinin resistant and sensitive parasites using 'Omics approaches.

| Publication | Level | No. Genes Used |
|---|---|---|
| Demas, et al. 2018 PNAS | Genome | 7 |
| Rocamora, et al. 2018 PloS Pathogens | Genome; Transcript | 47 |
| Mok, et al. 2021 Nature Comm. | Transcript | 80 |
| Siddiqui, et al. 2017 J. Infectious Disease | Protein | 12 |
| Witkowksi, et al. 2010 AAC. | Transcript | 9 |
| | **Total** | **154** |
| | **Unique Genes** | **151** |

| Gene ID | Publication |
|---|---|
| PF3D7_1454700 | Rocamora, et al. 2018 PloS Path |
| PF3D7_1457200 | Rocamora, et al. 2018 PloS Path |
| PF3D7_1457000 | Rocamora, et al. 2018 PloS Path |
| PF3D7_0704800 | Rocamora, et al. 2018 PloS Path |
| PF3D7_0730300 | Rocamora, et al. 2018 PloS Path |
| PF3D7_1115700 | Rocamora, et al. 2018 PloS Path |
| PF3D7_1141800 | Rocamora, et al. 2018 PloS Path |
| PF3D7_1427100 | Rocamora, et al. 2018 PloS Path |
| PF3D7_0420300 | Rocamora, et al. 2018 PloS Path |
| PF3D7_0528300 | Rocamora, et al. 2018 PloS Path |
| PF3D7_0617100 | Rocamora, et al. 2018 PloS Path |
| PF3D7_0810600 | Rocamora, et al. 2018 PloS Path |
| PF3D7_1230000 | Rocamora, et al. 2018 PloS Path |
| PF3D7_1368400 | Rocamora, et al. 2018 PloS Path |
| PF3D7_1478100 | Rocamora, et al. 2018 PloS Path |
| PF3D7_1200800 | Rocamora, et al. 2018 PloS Path |
| PF3D7_0702300 | Rocamora, et al. 2018 PloS Path |
| PF3D7_1030100 | Rocamora, et al. 2018 PloS Path |
| PF3D7_1300300 | Rocamora, et al. 2018 PloS Path |
| PF3D7_1479000 | Rocamora, et al. 2018 PloS Path |
| PF3D7_0933500 | Rocamora, et al. 2018 PloS Path |
| PF3D7_0812500 | Rocamora, et al. 2018 PloS Path |
| PF3D7_0113400 | Rocamora, et al. 2018 PloS Path |
| PF3D7_1110400 | Rocamora, et al. 2018 PloS Path |
| PF3D7_1116800 | Rocamora, et al. 2018 PloS Path |
| PF3D7_1235300 | Rocamora, et al. 2018 PloS Path |
| PF3D7_1351000 | Rocamora, et al. 2018 PloS Path |
| PF3D7_0609900 | Rocamora, et al. 2018 PloS Path |
| PF3D7_1406200 | Rocamora, et al. 2018 PloS Path |

| | |
|---|---|
| PF3D7_0400400 | Rocamora, et al. 2018 PloS Path |
| PF3D7_0600200 | Rocamora, et al. 2018 PloS Path |
| PF3D7_1001200 | Rocamora, et al. 2018 PloS Path |
| PF3D7_0400500 | Rocamora, et al. 2018 PloS Path |
| PF3D7_0712900 | Rocamora, et al. 2018 PloS Path |
| PF3D7_1139100 | Rocamora, et al. 2018 PloS Path |
| PF3D7_0201600 | Rocamora, et al. 2018 PloS Path |
| PF3D7_0302300 | Rocamora, et al. 2018 PloS Path |
| PF3D7_0600400 | Rocamora, et al. 2018 PloS Path |
| PF3D7_1208400 | Rocamora, et al. 2018 PloS Path |
| PF3D7_1470800 | Rocamora, et al. 2018 PloS Path |
| PF3D7_1245500 | Rocamora, et al. 2018 PloS Path |
| PF3D7_0409100 | Rocamora, et al. 2018 PloS Path |
| PF3D7_0617400 | Rocamora, et al. 2018 PloS Path |
| PF3D7_0918300 | Rocamora, et al. 2018 PloS Path |
| PF3D7_0906400 | Rocamora, et al. 2018 PloS Path |
| PF3D7_0107900 | Rocamora, et al. 2018 PloS Path |
| PF3D7_0315000 | Rocamora, et al. 2018 PloS Path |
| PF3D7_0701900 | Mok, et al. 2021 Nat Comm |
| PF3D7_1252600 | Mok, et al. 2021 Nat Comm |
| PF3D7_0702100 | Mok, et al. 2021 Nat Comm |
| PF3D7_1341700 | Mok, et al. 2021 Nat Comm |
| PF3D7_1343700 | Mok, et al. 2021 Nat Comm |
| PF3D7_1218500 | Mok, et al. 2021 Nat Comm |
| PF3D7_1252900 | Mok, et al. 2021 Nat Comm |
| PF3D7_0102200 | Mok, et al. 2021 Nat Comm |
| PF3D7_0401900 | Mok, et al. 2021 Nat Comm |
| PF3D7_1352900 | Mok, et al. 2021 Nat Comm |
| PF3D7_1002100 | Mok, et al. 2021 Nat Comm |
| PF3D7_0933000 | Mok, et al. 2021 Nat Comm |
| PF3D7_1108000 | Mok, et al. 2021 Nat Comm |
| PF3D7_1461600 | Mok, et al. 2021 Nat Comm |
| PF3D7_1006800 | Mok, et al. 2021 Nat Comm |
| PF3D7_0612900 | Mok, et al. 2021 Nat Comm |
| PF3D7_0901900 | Mok, et al. 2021 Nat Comm |
| PF3D7_1000800 | Mok, et al. 2021 Nat Comm |
| PF3D7_1361900 | Mok, et al. 2021 Nat Comm |
| PF3D7_0317200 | Mok, et al. 2021 Nat Comm |
| PF3D7_1304100 | Mok, et al. 2021 Nat Comm |
| PF3D7_1320100 | Mok, et al. 2021 Nat Comm |
| PF3D7_0811600 | Mok, et al. 2021 Nat Comm |
| PF3D7_0315900 | Mok, et al. 2021 Nat Comm |
| PF3D7_1246200 | Mok, et al. 2021 Nat Comm |

| | |
|---|---|
| PF3D7_0935800 | Mok, et al. 2021 Nat Comm |
| PF3D7_1223100 | Mok, et al. 2021 Nat Comm |
| PF3D7_1252100 | Mok, et al. 2021 Nat Comm |
| PF3D7_0104200 | Mok, et al. 2021 Nat Comm |
| PF3D7_1128900 | Mok, et al. 2021 Nat Comm |
| PF3D7_1467900 | Mok, et al. 2021 Nat Comm |
| PF3D7_1145200 | Mok, et al. 2021 Nat Comm |
| PF3D7_1335100 | Mok, et al. 2021 Nat Comm |
| PF3D7_0831600 | Mok, et al. 2021 Nat Comm |
| PF3D7_0722200 | Mok, et al. 2021 Nat Comm |
| PF3D7_1140400 | Mok, et al. 2021 Nat Comm |
| PF3D7_1206300 | Mok, et al. 2021 Nat Comm |
| PF3D7_1452000 | Mok, et al. 2021 Nat Comm |
| PF3D7_0810300 | Mok, et al. 2021 Nat Comm |
| PF3D7_0419700 | Mok, et al. 2021 Nat Comm |
| PF3D7_1021700 | Mok, et al. 2021 Nat Comm |
| PF3D7_0628100 | Mok, et al. 2021 Nat Comm |
| PF3D7_0905500 | Mok, et al. 2021 Nat Comm |
| PF3D7_0802600 | Mok, et al. 2021 Nat Comm |
| PF3D7_1022500 | Mok, et al. 2021 Nat Comm |
| PF3D7_0817600 | Mok, et al. 2021 Nat Comm |
| PF3D7_1035300 | Mok, et al. 2021 Nat Comm |
| PF3D7_1035900 | Mok, et al. 2021 Nat Comm |
| PF3D7_1104900 | Mok, et al. 2021 Nat Comm |
| PF3D7_1028700 | Mok, et al. 2021 Nat Comm |
| PF3D7_0614700 | Mok, et al. 2021 Nat Comm |
| PF3D7_1246400 | Mok, et al. 2021 Nat Comm |
| PF3D7_1229800 | Mok, et al. 2021 Nat Comm |
| PF3D7_1344300 | Mok, et al. 2021 Nat Comm |
| PF3D7_0407900 | Mok, et al. 2021 Nat Comm |
| PF3D7_1133400 | Mok, et al. 2021 Nat Comm |
| PF3D7_0822900 | Mok, et al. 2021 Nat Comm |
| PF3D7_0210600 | Mok, et al. 2021 Nat Comm |
| PF3D7_1036000 | Mok, et al. 2021 Nat Comm |
| PF3D7_1468400 | Mok, et al. 2021 Nat Comm |
| PF3D7_1243700 | Mok, et al. 2021 Nat Comm |
| PF3D7_0308300 | Mok, et al. 2021 Nat Comm |
| PF3D7_1251200 | Mok, et al. 2021 Nat Comm |
| PF3D7_0821400 | Mok, et al. 2021 Nat Comm |
| PF3D7_1310700 | Mok, et al. 2021 Nat Comm |
| PF3D7_0515400 | Mok, et al. 2021 Nat Comm |
| PF3D7_1036500 | Mok, et al. 2021 Nat Comm |
| PF3D7_0424100 | Mok, et al. 2021 Nat Comm |

| | |
|---|---|
| PF3D7_1125700 | Mok, et al. 2021 Nat Comm |
| PF3D7_0214600 | Mok, et al. 2021 Nat Comm |
| PF3D7_0828800 | Mok, et al. 2021 Nat Comm |
| PF3D7_1227700 | Mok, et al. 2021 Nat Comm |
| PF3D7_0402300 | Mok, et al. 2021 Nat Comm |
| PF3D7_1238800 | Mok, et al. 2021 Nat Comm |
| PF3D7_0731500 | Mok, et al. 2021 Nat Comm |
| PF3D7_0408000 | Mok, et al. 2021 Nat Comm |
| PF3D7_1230700 | Mok, et al. 2021 Nat Comm |
| PF3D7_1433500 | Mok, et al. 2021 Nat Comm |
| PF3D7_0410000 | Mok, et al. 2021 Nat Comm |
| PF3D7_1240100 | Mok, et al. 2021 Nat Comm |
| PF3D7_1251200 | Demas, et al. 2018 PNAS |
| PF3D7_1433800 | Demas, et al. 2018 PNAS |
| PF3D7_1126100 | Demas, et al. 2018 PNAS |
| PF3D7_0209600 | Demas, et al. 2018 PNAS |
| PF3D7_1121900 | Demas, et al. 2018 PNAS |
| PF3D7_1324300 | Demas, et al. 2018 PNAS |
| PF3D7_1422400 | Demas, et al. 2018 PNAS |
| PF3D7_1343700 | Siddiqui, et al. 2017 JID |
| PF3D7_0500800 | Siddiqui, et al. 2017 JID |
| PF3D7_1116700 | Siddiqui, et al. 2017 JID |
| PF3D7_0406200 | Siddiqui, et al. 2017 JID |
| PF3D7_0702500 | Siddiqui, et al. 2017 JID |
| PF3D7_1343700 | Siddiqui, et al. 2017 JID |
| PF3D7_1364800 | Siddiqui, et al. 2017 JID |
| PF3D7_0209800 | Siddiqui, et al. 2017 JID |
| PF3D7_1447000 | Siddiqui, et al. 2017 JID |
| PF3D7_0204500 | Siddiqui, et al. 2017 JID |
| PF3D7_1206200 | Siddiqui, et al. 2017 JID |
| PF3D7_0320700 | Siddiqui, et al. 2017 JID |
| PF3D7_0202000 | Wikowski, et al. 2010 AAC |
| PF3D7_1372500 | Wikowski, et al. 2010 AAC |
| PF3D7_1372600 | Wikowski, et al. 2010 AAC |
| PF3D7_0818900 | Wikowski, et al. 2010 AAC |
| PF3D7_1012400 | Wikowski, et al. 2010 AAC |
| PF3D7_0202100 | Wikowski, et al. 2010 AAC |
| PF3D7_0300900 | Wikowski, et al. 2010 AAC |
| PF3D7_0301700 | Wikowski, et al. 2010 AAC |
| PF3D7_0528400 | Wikowski, et al. 2010 AAC |
